# Validating Minimally Invasive Laser Doppler Flowmetry for Serial Bone Perfusion Measurements in Mice

**DOI:** 10.1101/708412

**Authors:** Nicholas J. Hanne, Elizabeth D. Easter, Jacqueline H. Cole

**Author notes:** These authors contributed equally to this work (Nicholas J. Hanne, Elizabeth D. Easter). **Corresponding Author:** Jacqueline H. Cole, Joint Department of Biomedical Engineering, University of North Carolina and North Carolina State University, 911 Oval Drive, Campus Box 7115, Raleigh, NC 27695-7115.

## Abstract

*In vivo* laser Doppler flowmetry (LDF) has previously been used to quantify blood perfusion accurately at a single timepoint in the murine tibial metaphysis. However, this procedure entailed substantial disruption to soft tissues overlying the bone and caused notable localized inflammation for several weeks after the procedure, impeding serial measurements in the same mouse. In this study, we tested a less invasive technique to measure perfusion in the tibia with LDF and validated that it can be used serially in the same mouse without causing inflammation or gait perturbations. Twenty 14-week-old C57Bl/6J mice were evenly divided into groups that either had daily treadmill exercise or remained sedentary. Within these activity groups, mice were evenly subdivided into groups that received LDF measurements either weekly or only once at the study endpoint. Bone perfusion was measured with LDF in the anteromedial region of the right tibial metaphysis. Serum concentrations of interleukin 6, incision site wound area, and interlimb coordination during gait were measured weekly for four weeks. Tibial perfusion did not differ significantly between exercise and sedentary groups within the weekly or endpoint-only LDF groups at any timepoint. Perfusion was significantly increased in the third week in the weekly LDF group relative to measurements in the second and fourth weeks. Ligation of the femoral artery caused consistent, rapid reductions in tibial perfusion, validating that LDF is sensitive to changes in tibial blood supply. Weekly LDF procedures did not adversely affect gait, as interlimb coordination during treadmill locomotion was similar between weekly and endpoint-only LDF groups at every timepoint. Images of the incision site show wound closure within one week, and serum concentrations of interleukin 6 were not significantly different between weekly and endpoint-only groups. Together, these findings demonstrate that our minimally invasive LDF technique can be used for serial *in vivo* measurements of intraosseous blood perfusion without inducing localized inflammation or negatively affecting gait patterns in mice.

**Highlights:**

- Modified, minimally invasive laser Doppler flowmetry (LDF) technique was validated for serial measures of tibial perfusion in mice.
- Weekly LDF procedures did not induce inflammation or alter gait patterns that could confound metrics of interest in bone studies.
- Ligation of the femoral artery confirmed the LDF technique measures functional perfusion within the bone.

## 1. Introduction

Vasculature within bone (*osteovasculature*) is an essential contributor to bone health, providing nutrients, oxygen, cells, and chemical signals and removing waste products [1,2]. Adequate vascular perfusion is required for bone development, adaptation in response to loading, and healing after fracture [2–4]. Evidence that vascular pathologies are associated with bone loss is growing. Aortic calcification is associated with decreased lumbar spine bone mineral density (BMD) and increased fracture risk in men and women within four years [5], and the incidence of cardiovascular disease increases with reduced BMD in the spine in white men, and hip, trochanter, and femoral neck in black women [6]. Osteoporosis is associated with reduced perfusion in the vertebrae for men [7] and in the femoral head for women [8], although the mechanisms responsible for bone loss in these individuals is unknown and needs further examination using animal studies. Although murine models are commonly used to determine the effect of pathologies on bone properties, measuring blood perfusion within mouse bone is complicated due to their small size. Current methods are either experimentally difficult (e.g., hydrogen washout [9]) or require the animal to be sacrificed (e.g., microspheres, radiolabels, polyoxometalates, barium sulfate, or Microfil^®^ [10–16]). Some methods can be performed *in vivo* but provide poor resolution not suitable for small bones: laser speckle imaging [17], laser Doppler perfusion imaging [18], contrast-enhanced magnetic resonance imaging (MRI) [19], contrast-enhanced positron emission tomography (PET) [19], and contrast-enhanced micro-computed tomography [20,21]. Endpoint and *ex vivo* measurements only provide a snapshot of vascular network function, missing the timing of vascular changes and any transient changes to vascular supply. A technique that could be used for longitudinal studies of bone perfusion *in vivo* would enable us to capture temporal changes in bone perfusion for individual subjects, thereby improving understanding of disease progression and intervention effectiveness.

First proposed as a tool to measure intraosseous perfusion by Nilsson *et al*. in 1980 [22], laser Doppler flowmetry (LDF) directs a monochromatic light source over a perfused tissue and measures backscattered light from fluid movement with a photodetector to provide a relative measure of blood perfusion. Perfusion is a functional measure of blood flow that is affected not only by the amount and velocity of red blood cells but also capillary density, vascular permeability, and flow direction [23,24]. LDF was first used to measure blood perfusion in the cancellous bone of pig mandibles by Hellem *et al*. in 1983 [25] and thereafter was rapidly adopted in orthopaedic clinics as an intraoperative tool to aid surgeons in identifying non-viable bone for debridement in patients with osteomyelitis, osteonecrosis of the femoral head, and lower limb traumatic injury [23]. LDF has also been used as an endpoint measure to compare relative intraosseous perfusion between groups in murine research studies [26,27]. Recently, it was validated as a tool to quantify perfusion in the mouse tibia, but the technique used in that study involved a relatively large incision that resulted in inflammation at the incision site up to three months after the procedure [28]. To monitor longitudinal changes in murine bone perfusion, a less invasive LDF procedure is needed that will not induce significant localized inflammation or limping during gait, which could have both biological and mechanical confounding effects on bone.

We developed a minimally invasive LDF procedure and have used it to measure changes in tibial perfusion in mice in response to diet-induced obesity, ischemic stroke, and treadmill exercise [29,30]. The objective of this study was to determine if this modified LDF procedure could be performed repeatedly in a longitudinal study without affecting bone perfusion, inducing inflammation, or altering limb coordination during locomotion, which could confound bone metrics of interest and perfusion changes associated with interventions. Developing new *in vivo* techniques to measure bone blood perfusion is a critical step needed to understand longitudinal changes to osteovasculature, which may contribute to bone loss occurring during progression of various clinical pathologies.

## 2. Materials and Methods

### 2.1 Study design

The protocol for this study was approved by the Institutional Animal Care and Use Committee at North Carolina State University. Eighteen 14-week-old male C57Bl/6J mice (The Jackson Laboratory, Bar Harbor, ME) were acclimated to the animal facility for one week. They were group-housed (4 per cage) on a 12-hour light/12-hour dark diurnal cycle and provided chow and water *ad libitum*. The mice were randomly assigned to four groups (Fig. 1) based on LDF procedure frequency (weekly, endpoint) and exercise regimen (sedentary, exercise): weekly sedentary (n = 5), weekly exercise (n = 5), endpoint sedentary (n = 4), and endpoint exercise (n = 4). Exercise groups were acclimated to treadmill exercise (Exer-3/6, Columbus Instruments, Columbus, OH) by increasing exercise intensity for 2 days prior to the start of the study (Day 1: 5 m/min for 10 min, 9 m/min for 10 min, and 12 m/min for 10 min; Day 2: 5 m/min for 5 min, 9 m/min for 5 min, and 12 m/min for 20 min). During the study, exercise groups performed daily exercise for 4 weeks (30 min/day, 5 days/week, 12 m/min, 5° incline), while sedentary groups were placed on a stationary treadmill for the same amount of time to equalize handling among the groups.

**Figure 1.**
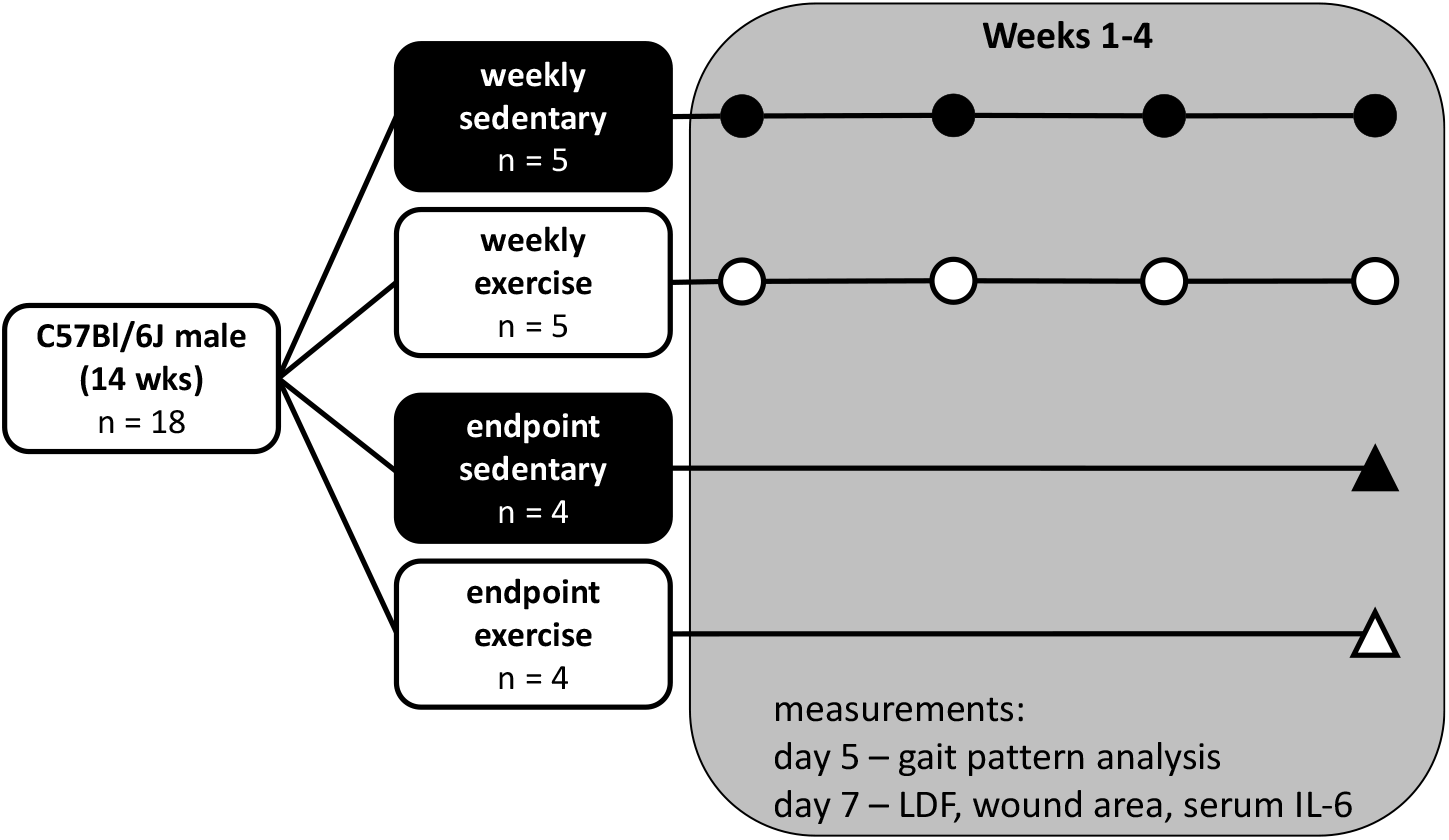
Experimental design. Symbols indicate weeks in which tibial perfusion was measured with LDF. Wound area, serum interleukin 6 (IL-6), and gait patterns were assessed weekly.

Laser Doppler flowmetry was used to measure intraosseous blood perfusion in the right tibial metaphysis, performed either weekly for 4 weeks (weekly groups) or at a single timepoint at the end of the study (endpoint groups). For the weekly LDF groups, starting in Week 2, images of the incision site were taken during the LDF procedure prior to making an incision to assess the wound from the previous week. Blood samples were collected from the submandibular vein of all mice under anesthesia (at the end of the LDF procedure for the weekly groups). Blood samples were centrifuged at 2,000 x g for 10 min, and the isolated serum was stored at −80°C until analysis with an enzyme-linked immunosorbent assay (ELISA). At five days after each of the first three LDF procedures, gait patterns were assessed using high-speed video. Immediately following the last LDF procedure and serum collection, mice were euthanized using CO2 asphyxiation followed by cervical dislocation.

### 2.2 Tibial perfusion

All mice were fasted for 6-8 hours before each LDF procedure. Anesthesia was induced and maintained with isoflurane (2%) in pure oxygen throughout the procedure (about 15 minutes). After anesthesia induction, the fur over the right knee was shaved, mice were placed supine on a heated pad, and the right leg was taped to the surgical platform. A 2-5-mm long incision was made over the anteromedial surface of the right proximal tibial metaphysis, the bone was exposed, and a small region of the periosteum was scraped away. LDF measurements were recorded using an LDF monitor with a 785-nm light source (MoorVMS-LDF, Moor Instruments Ltd, Axminster, UK), and a 3 kHz lowpass filter selected. A VP4 Needle Probe (0.8 mm outer diameter, 0.25 mm fiber separation) was placed directly on the exposed bone surface (Fig. 2) and held in place using a micromanipulator (MM3-ALL, World Precision Instruments, Sarasota, FL) to reduce signal noise from probe movement. Each weekly measurement was composed of the weighted mean and standard deviation of three 30-second readings, with repositioning of the probe between readings. The incisions were closed using VetBond™ tissue glue (3M Company, St. Paul, MN) and covered with triple antibiotic cream.

**Figure 2.**
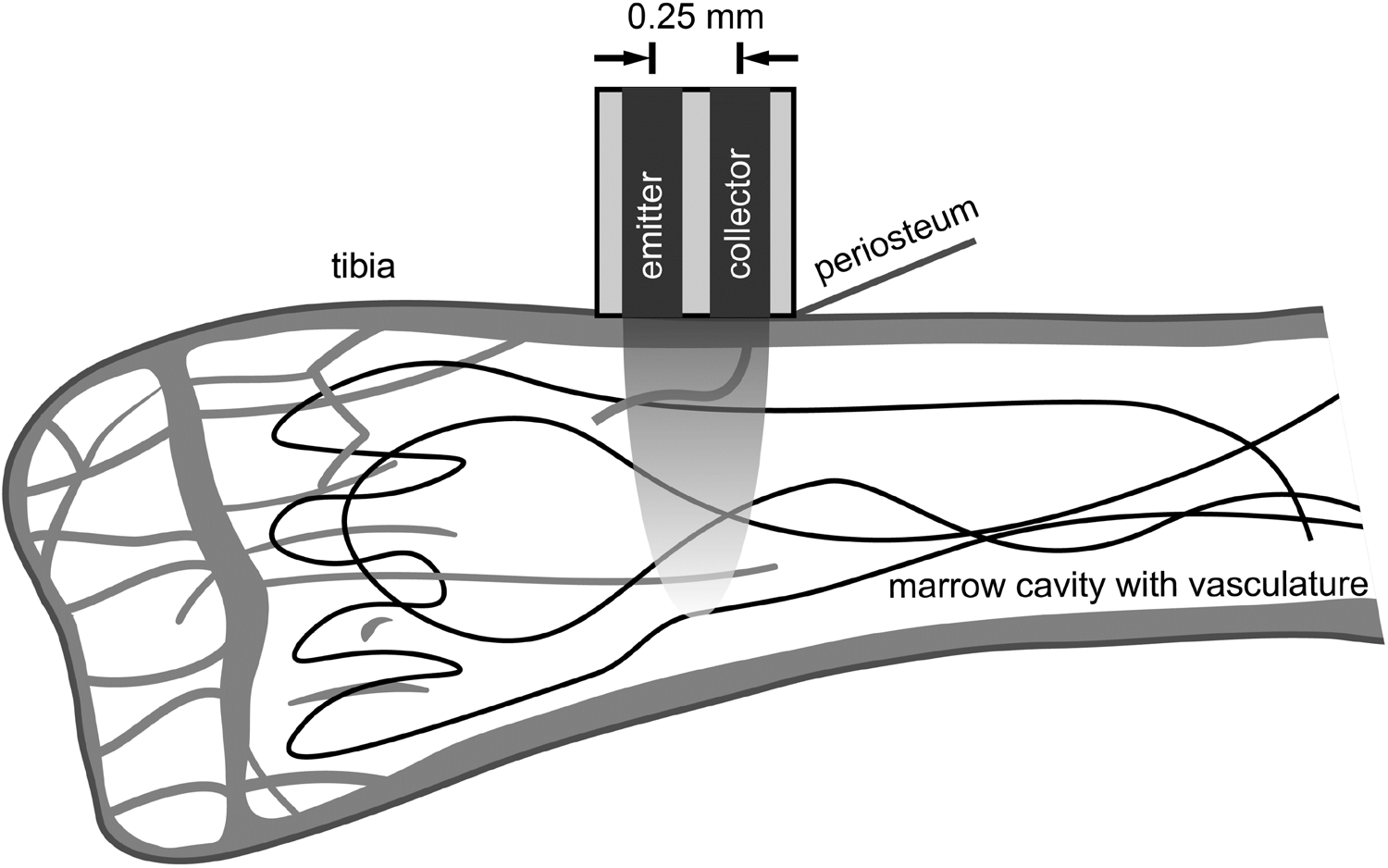
Schematic of LDF setup for bone perfusion measurements in the proximal tibia. The probe, placed on the tibial surface, emitted 785-nm light that scattered through a region of underlying tissue (represented by parabolic shading) and experienced Doppler shifts, some of which was scattered back to the collection probe, where it was measured.

### 2.3 Femoral ligation validation

At the end of the study, just prior to euthanasia, additional LDF measurements were performed in a subset of mice (n=12) during arterial ligation to confirm the association of LDF perfusion measurements with changes in blood supply to the bone. While still anesthetized during the final LDF procedure, the skin incision over the proximal tibia was extended to expose the entire inner thigh and the femoral artery. The LDF probe was again positioned over the tibial metaphysis, and a suture was tied around the femoral artery but not tightened. A 30-second baseline measurement was taken, the suture was tightened to ligate the artery, and another 30-second measurement was recorded. The reduction in tibial perfusion was calculated as the ratio of the ligated measurement to the baseline measurement, expressed as a percent.

### 2.4 Wound area

As mentioned above, for the weekly LDF groups, pictures were taken of the incision wounds immediately prior to each LDF procedure to assess localized inflammation and healing at the wound site from the procedure performed in the preceding week. Wound area was calculated by tracing the edge of the wound in ImageJ (version 1.51k, National Institutes of Health, Bethesda, MD). The wound was considered closed if no moist granulation tissue was visible and the wound was covered with new epithelium [31].

### 2.5 Serum concentration of interleukin 6

Systemic inflammation was examined by quantifying serum concentrations of the proinflammatory marker interleukin 6 (IL-6) with an ELISA (IL-6 Mouse ELISA kit, KMC0061, Invitrogen, Carlsbad, CA). Samples were prepared according to the manufacturer’s instructions and measured using a plate reader (Synergy H1, BioTek Instruments, Inc., Winooski, VT). Due to limited serum, some samples were diluted 2-4 times to allow samples to be run in duplicate.

### 2.6 Gait pattern analysis

The effect of the LDF procedures on interlimb coordination was examined weekly in all mice, five days after each procedure day, because limping or other gait asymmetries could alter the strain experienced by hindlimb bones and confound bone outcome measures [32]. During a short treadmill session (12 m/min for 60 sec), high-speed video was collected in the sagittal plane at 240 frames per second (HERO4, GoPro, Inc., San Mateo, CA). Gait was analyzed using Kinovea (version 0.8, Kinovea Open Source Project) to quantify duty cycle for both hindlimbs and phase dispersions for ipsilateral, diagonal, and contralateral limbs with relation to the LDF limb (right hindlimb) (Fig. 3) [33–35]. Duty cycle for a given limb is the ratio of the time that limb is on the ground (measured from paw strike to lift off) to the total time of an entire gait cycle (measured from paw strike to paw strike). Phase dispersion between two limbs is a measure of the time between paw strikes for those two limbs within a gait cycle. Five consecutive gait cycles were analyzed for each treadmill session. Duty cycle and phase dispersion were averaged over the five gait cycles at each timepoint.

**Figure 3.**
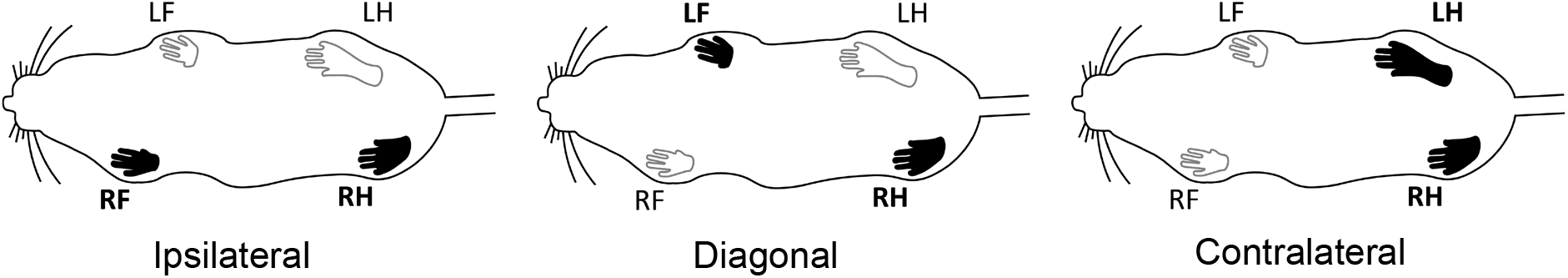
Schematic of the ipsilateral (RF-RH), diagonal (LF-RH), and contralateral (LH-RH) phase dispersions used for gait analysis. Relevant limbs relative to the LDF limb (RH) shaded in black. L=left, R=right, F=forelimb, H=hindlimb.

### 2.7 Statistical analysis

All data analyses were performed using SAS (SAS University Edition v. 9.4, SAS Institute Inc., Cary, NC) with a significance level of 0.05. Models were chosen to answer five questions: 1) Does performing weekly LDF procedures affect bone perfusion? To answer this question, LDF data from the final timepoint (Week 4) were compared across LDF frequency (weekly, endpoint) and exercise regimen (sedentary, exercise) using a two-way ANOVA with interaction. Tukey’s post-hoc tests were used to compare group means. 2) Does exercise affect bone perfusion? For this question, LDF data for the weekly group were compared across exercise regimen and timepoint (Weeks 1-4) to examine differences between exercise groups within each timepoint (i.e., sedentary vs. exercise at Week 1). A mixed effects general linear model (procedure MIXED) with interaction was used, with exercise group as a fixed factor and timepoint as a repeated factor. The covariance matrix was modeled using compound symmetry. Exercise effect differences were calculated based on least squares means (LSM) with Tukey-Kramer adjustments for multiple comparisons. For the endpoint-only group (Week 4 data), the effect of exercise was examined with one-way ANOVA. 3) Do weekly LDF procedures alter exercise effects on bone perfusion? Effect differences between timepoints (i.e., Week 1 vs. Week 2) were evaluated in the same mixed effects model using LSM with Tukey-Kramer adjustments for multiple comparisons. 4) Do LDF readings directly correspond to changes in blood supply, assessed by femoral artery ligation? LDF data before and after the femoral artery was ligated were compared using a paired t-test. 5) Do weekly LDF procedures alter gait and interlimb coordination? This question was addressed by comparing gait parameters between LDF groups within exercise groups and timepoints (i.e., weekly sedentary vs. endpoint sedentary at Week 1). Four gait parameters were compared across LDF groups, exercise groups, and timepoints (Weeks 2-4) with mixed effects linear models (procedure MIXED) with interaction, where LDF frequency and exercise regimen were fixed factors and timepoint was a repeated measure. The covariance matrices were modeled using either compound symmetry (duty cycle, contralateral phase dispersion) or a first-order autoregressive (diagonal and ipsilateral phase dispersion), based on which covariance matrix yielded a lower corrected Akaike’s Information Criterion. Effect differences were calculated based on LSM with Tukey-Kramer adjustments for multiple comparisons. All data are presented as mean ± standard deviation, except LDF and gait data, which are presented as LSM ± 95% confidence interval.

## 3. Results

### 3.1 Tibial perfusion

At Week 4, bone perfusion was similar between the weekly and endpoint-only groups (p = 0.92), showing that our modified LDF procedure can be performed weekly without affecting perfusion, the primary outcome of interest (Fig. 4). With weekly LDF procedures, treadmill exercise did not affect perfusion measurements, compared to the sedentary group, at any timepoint (p = 0.11 Week 1, p = 0.64 Week 2, p = 0.76 Week 3, and p = 0.78 Week 4). Similarly, with endpoint-only LDF procedures, perfusion also did not differ significantly between exercise and sedentary groups at Week 4 (p = 0.49), suggesting weekly LDF procedures did not mask an exercise effect on perfusion. Perfusion was higher in Week 3 compared to Weeks 2 and 4, indicating that the technique is sensitive to transient perfusion changes regardless of exercise.

**Figure 4.**
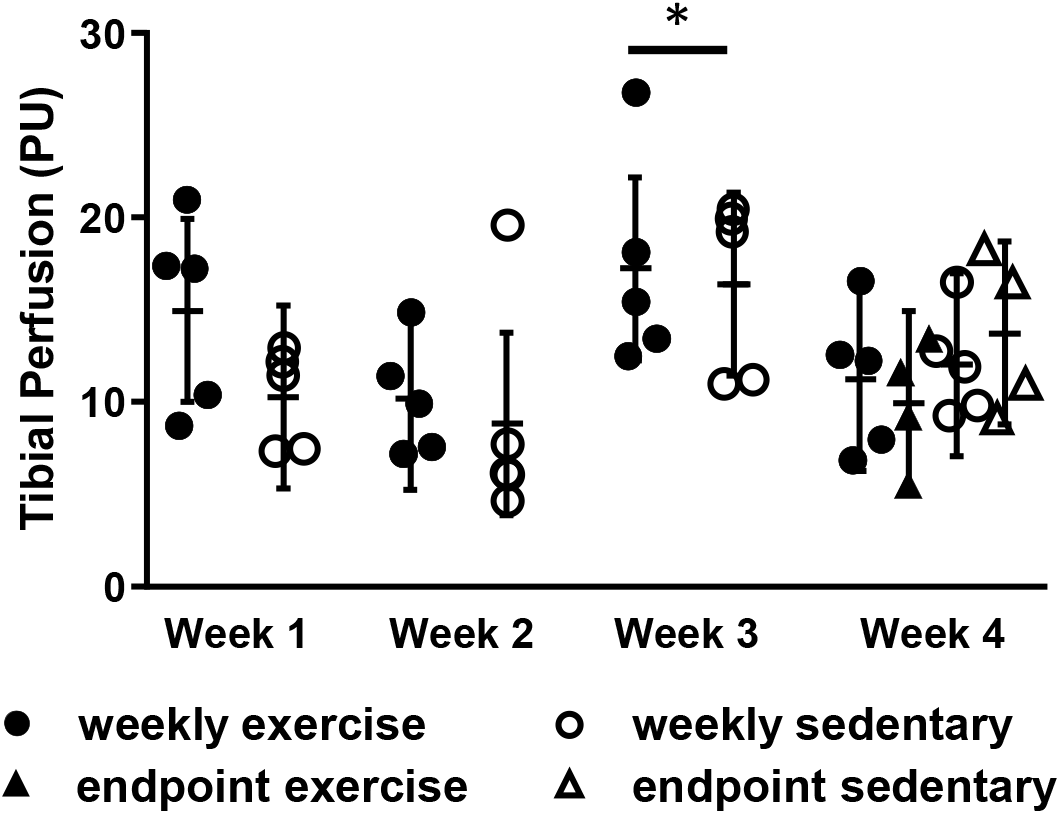
At Week 4, tibial perfusion was similar between weekly and endpoint-only groups. Sedentary and exercise groups had similar perfusion at every timepoint. LDF was sensitive to transient perfusion changes, measuring higher in Week 3 compared to Week 2 and Week 4. PU = perfusion units. Data presented as least squares mean ± 95% confidence interval. *p < 0.05 vs. Weeks 2 and 4.

### 3.2 Femoral ligation validation

After the femoral artery was ligated, perfusion in the tibia dropped rapidly from 15.3 ± 9.2 perfusion units (PU) to 3.8 ± 1.4 PU within 30 seconds (31 ± 19% of the baseline perfusion, p = 0.004) (Fig. 5), validating that the LDF perfusion measurements are directly associated with blood supply within the bone. Two measurements were not included in the analysis, because the LDF probe slipped off the tibia when the ligature was tightened.

**Figure 5.**
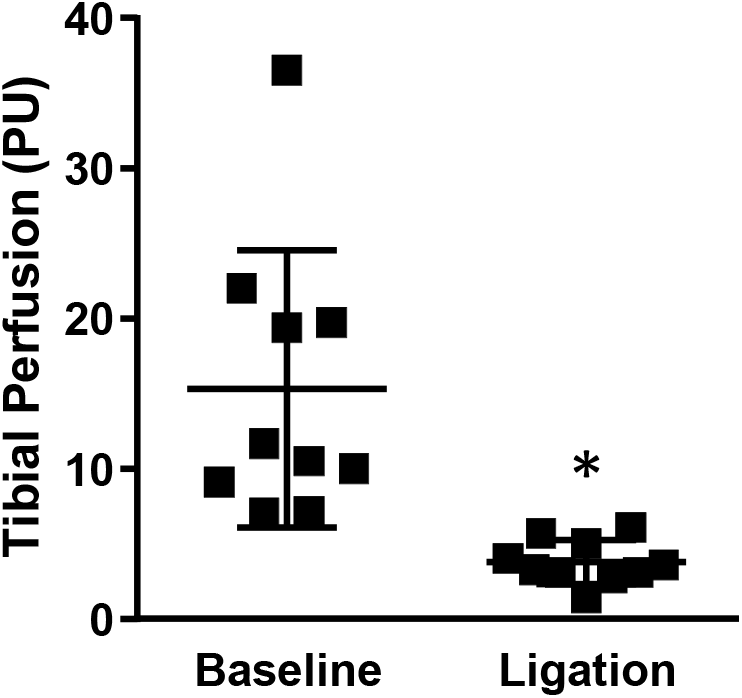
After ligation of the femoral artery, tibial perfusion dropped to 31 ± 19% of the baseline value. PU = perfusion units. *p < 0.05 vs. baseline.

### 3.3 Wound area

Wound images taken one week after each weekly LDF procedure showed minimal signs of inflammation for all but one incision for both exercise and sedentary mice, with either full closure (Fig. 6A) or a small, dry scab (Fig. 6B) resulting in zero wound area recorded. Wet granulation tissue was observed in only one incision site, for a sedentary mouse in Week 2 (Fig. 6C, wound area = 0.39 mm^2^), and the wound was healed within the subsequent week.

**Figure 6.**
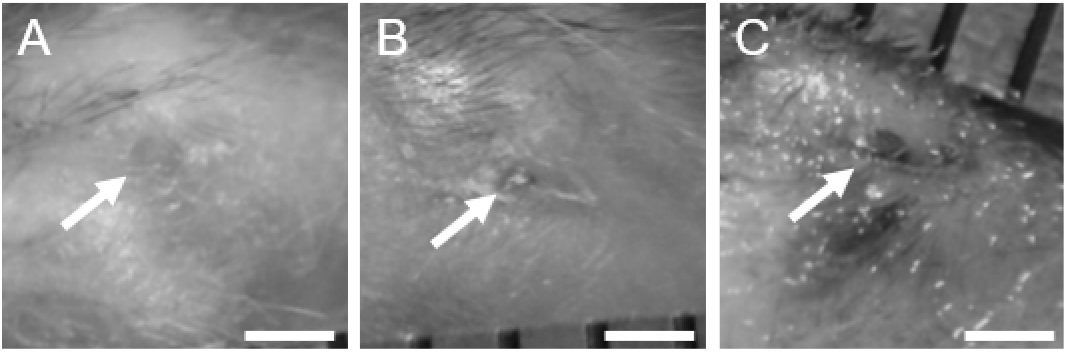
Example images of the incision wound site (arrows) from the weekly LDF group taken one week following the procedure. Incisions were either A) fully closed or B) closed with small, dry granulation tissue. C) Only one incision did not fully heal, but it was healed by the following week. Scale bars are 1 mm.

### 3.4 Serum concentration of IL-6

Circulating levels of proinflammatory marker IL-6 were below the detectable limit for all animals at each timepoint, except one mouse in the endpoint sedentary group at Week 4 (52.1 pg/mL). Since the lower threshold of the test is 7.8 pg/mL, and serum samples were diluted up to four times to allow for duplicate measurements, serum levels of IL-6 were below 31.2 pg/mL. These levels agree with normal physiologic concentrations of IL-6, which are below 100 pg/mL in C57Bl/6J mice [36,37], suggesting the mice in our study experienced little to no systemic inflammation in response to the weekly LDF procedures. Pathologic inflammation can increase IL-6 levels up to 200-1,000 pg/mL [36,37]

### 3.5 Gait pattern analysis

Weekly LDF procedures did not affect gait parameters during treadmill locomotion. Limb coordination did not differ between weekly and endpoint-only groups at any timepoint for either sedentary or exercise groups (Fig. 7). Sedentary groups did have small alterations in gait parameters in Week 2 compared to exercise groups (hindlimb duty cycle and diagonal and contralateral phase dispersion), possibly indicating slight discomfort or unfamiliarity with treadmill locomotion. Diagonal phase dispersion was lower in exercise mice compared to sedentary during Week 4 (8.6% lower, p = 0.012).

**Figure 7.**
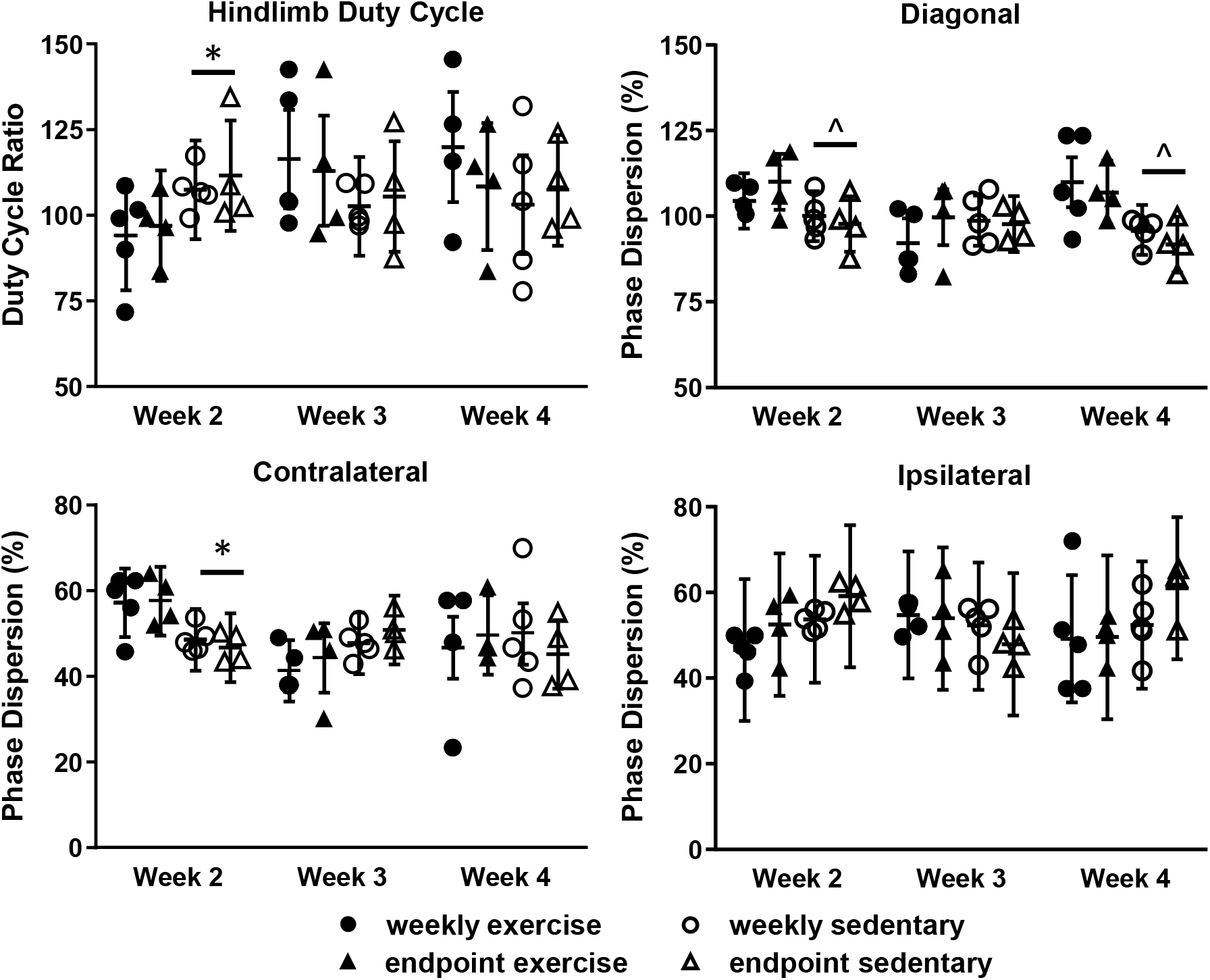
Treadmill locomotion patterns involving the LDF affected limb (right hindlimb). Gait patterns were not significantly different between weekly and endpoint-only groups at any timepoint for hindlimb duty cycle ratio (A), diagonal phase dispersion (B), contralateral phase dispersion (C), or ipsilateral phase dispersion (D). Diagonal and contralateral phase dispersion were lower in exercise than sedentary groups at some timepoints. Data are presented as least squares mean ± 95% confidence interval. *p < 0.05 vs. sedentary. ^ p = 0.066 vs. sedentary.

## 4. Discussion

Our minimally invasive laser Doppler flowmetry technique measured *in vivo* intraosseous perfusion in the tibia weekly without inducing localized or systemic inflammation. LDF measures of tibial perfusion were similar between groups that received weekly procedures and groups that received only an endpoint procedure, indicating that the procedure itself did not impact measurements of intraosseous perfusion. A previous study demonstrated that LDF could be used to quantify blood perfusion in murine tibiae but noted that signs of inflammation were observed at the incision site up to three months following the procedure and suggested that it be used as an endpoint-only measure to avoid influencing bone outcomes [28]. In this study, we limited the invasiveness of the procedure by reducing the incision size and preserving muscle tissue, thereby enabling repeated measurements to be performed without causing chronic increases in proinflammatory marker IL-6. Furthermore, all but one incision from the procedures were fully closed within the one-week period before the next procedure, and no visual signs of inflammation were observed throughout the study. The fast healing times observed in the small incisions from this procedure (approximately 1 mm) are consistent with wound closure studies, where even large (9-28 mm^2^), unclosed dermal biopsies heal in 7-14 days [38].

The femoral ligation test confirmed that LDF measures are sensitive to changes in blood supply and thus are directly related to perfusion within the underlying cortical bone and marrow space. We demonstrated that femoral artery ligation reliably decreased the measured perfusion to about 3.8 PU (31% of baseline) within 30 seconds in all mice. Cortical thickness in the tibia affects the depth into the marrow space that LDF can measure. Another study quantified this relationship in the murine tibia and found that small variations in cortical thickness had minimal effect on LDF perfusion measurements [28], suggesting that LDF can be used to track longitudinal changes in bone perfusion. Because cortical thickness in the tibial diaphysis changes by ~10% from skeletal maturity at 16 weeks of age to 52 weeks of age [39], and can differ by sex [40], longitudinal LDF measurements should only be performed in age- and sex-matched subjects. Finally, we found no interaction effect of daily treadmill exercise on LDF readings. Taken together, these results suggest that the minimally invasive LDF procedure validated in our study can be used to monitor and compare blood perfusion longitudinally in murine studies involving exercise therapy. This procedure can be used to track changes to osteovascular function, which is known to play an important role in bone development, remodeling, and repair [41,42], yet remains under-studied.

In addition to providing aerobic exercise, treadmill activity also mechanically loads the bones and increases the functional strain experienced by the bones [43–45]. Even slight changes to functional strain can affect osteogenesis [32,46] and angiogenesis [15,47]. Since other studies have shown that changes to gait kinematics and limb patterning affect bone strain [44,48], we were concerned the LDF procedure could affect locomotion patterns and confound exercise effects by altering functional strain. We found no differences in duty cycle or interlimb coordination between weekly and endpoint-only groups at any timepoint, indicating that weekly LDF measurements do not alter gait patterns (and thus functional strain) during treadmill exercise.

Although not the main focus of this study, aerobic treadmill exercise was anticipated to cause increased tibial perfusion over time due to vascular growth and adaptation, as we have previously found in studies using the same exercise regimen and LDF technique [29,30]. Perfusion is a functional measure of not only the amount and direction of blood flow but also vascular permeability and capillary density [24]. Treadmill exercise may be affecting the size, number, or cellular makeup of the vasculature without causing changes in perfusion. Rats that performed a similar treadmill routine for two weeks had a 19% increase in the number but not total area of blood vessels in the proximal tibial metaphysis compared to the sedentary group [15]. A more rigorous aerobic exercise intervention, such as free access to running wheels where mice will run 4-10 km daily [49,50], may have a larger and more detectible effect on perfusion. Nevertheless, our study confirmed that weekly LDF procedures will not confound perfusion measurements in future exercise studies.

Stress, like inflammation and aerobic exercise, can also affect vascular function. Neuropeptide Y, which is expressed during stress response, has been shown to be both angiogenic and vasoconstrictive, which could increase blood pressure and the amount of vasculature in bone [51]. Although we found no detectible increases in IL-6 or visible signs of inflammation at the incision site, both sedentary and exercise mice had a significant increase in tibial perfusion in the third week that was resolved by the fourth week. Stress may have played a role in increasing perfusion in the third week; all mice were handled daily for either treadmill exercise or the sedentary treadmill activity and had blood drawn weekly which could induce a stress response. The increased perfusion lasted only one week and was present in both exercise and sedentary mice.

This study had several limitations that warrant attention in future studies. A primary concern of this technique is the removal of a small area of the periosteum, a highly vascularized tissue that contains osteoblast precursor cells [52]. LDF measures of intracortical perfusion in the tibiae of juvenile ewes dropped by 25% immediately following the removal of the periosteum from the medial aspect [53]. Although the amount of periosteum removed in our procedure is small (about the size of our probe, 0.5 mm^2^ area), the effects of periosteal removal were not examined and may affect bone tissue function. The effects of weekly LDF procedures on bone remodeling and homeostasis were not examined in this study. This study did not compare LDF results to other promising emerging techniques for examining osteovasculature *in vivo*. Several new higher resolution PET scans can be used in rodent bones [54], and emerging MRI techniques (e.g., blood oxygen level-dependent MRI and intravoxel incoherent motion MRI [24]) greatly improve resolution and do not require contrast agents, but these techniques remain prohibitively expensive.

## 5. Conclusions

Weekly LDF procedures performed over four weeks did not induce measurable signs of inflammation or significantly alter gait patterns during treadmill exercise. Unlike other existing methods used for measuring the vascular network in bone, this procedure can be performed *in vivo*, is repeatable without confounding study controls, is relatively simple to perform, and is inexpensive. Monitoring intraosseous perfusion serially with LDF provides a functional measure of blood flow, enabling researchers to track changes to the osteovasculature noninvasively during disease progression and interventions.

## Acknowledgments

This work was supported by the Eunice Kennedy Shriver National Institute of Child Health and Human Development (NICHD) of the National Institutes of Health (NIH) under award number K12HD073945. Additional support was provided by the Office of Undergraduate Research at North Carolina State University.

## Abbreviations

LDF: laser Doppler flowmetry

## References

[1] A.M. Parfitt, The mechanism of coupling: a role for the vasculature, Bone. 26 (2000) 319–323. doi:10.1016/S8756-3282(00)80937-0.

[2] J. Trueta, The Role of the Vessels in Osteogenesis, Bone Joint J. 45-B (1963) 402–418.

[3] R.E. Tomlinson, J.A. McKenzie, A.H. Schmieder, G.R. Wohl, G.M. Lanza, M.J. Silva, Angiogenesis is Required for Stress Fracture Healing in Rats, Bone. 52 (2013) 212–219. doi:10.1016/j.bone.2012.09.035.

[4] A.L. Wallace, E.R.C. Draper, R.K. Strachan, I.D. Mccarthy, S.P.F. Hughes, The effect of devascularisation upon early bone healing in dynamic external fixation, Bone Joint J. 73 (1991) 819–825.

[5] M. Naves, M. Rodríguez-García, J.B. Díaz-López, C. Gómez-Alonso, J.B. Cannata-Andía, Progression of vascular calcifications is associated with greater bone loss and increased bone fractures, Osteoporos Int. 19 (2008) 1161–1166. doi:10.1007/s00198-007-0539-1.

[6] G.N. Farhat, A.B. Newman, K. Sutton-Tyrrell, K.A. Matthews, R. Boudreau, A.V. Schwartz, T. Harris, F. Tylavsky, M. Visser, J.A. Cauley, for the H.A. Study, The association of bone mineral density measures with incident cardiovascular disease in older adults, Osteoporos Int. 18 (2007) 999–1008. doi:10.1007/s00198-007-0338-8.

[7] J.F. Griffith, D.K.W. Yeung, G.E. Antonio, F.K.H. Lee, A.W.L. Hong, S.Y.S. Wong, E.M.C. Lau, P.C. Leung, Vertebral Bone Mineral Density, Marrow Perfusion, and Fat Content in Healthy Men and Men with Osteoporosis: Dynamic Contrast-enhanced MR Imaging and MR Spectroscopy, Radiology. 236 (2005) 945–951. doi:10.1148/radiol.2363041425.

[8] J.F. Griffith, D.K. Yeung, P.H. Tsang, K.C. Choi, T.C. Kwok, A.T. Ahuja, K.S. Leung, P.C. Leung, Compromised Bone Marrow Perfusion in Osteoporosis, J Bone Miner Res. 23 (2008) 1068–1075. doi:10.1359/jbmr.080233.

[9] L.A. Whiteside, P.A. Lesker, D.J. Simmons, Measurement of Regional Bone and Bone Marrow Blood Flow in the Rabbit Using the Hydrogen Washout Technique, Clin. Orthop. Relat. Res. 122 (1977) 340–346.

[10] M.A. Serrat, Measuring bone blood supply in mice using fluorescent microspheres, Nat. Protoc. 4 (2009) 1779–58. doi:http://dx.doi.org/10.1038/nprot.2009.190.

[11] O. Grundnes, O. Reikerås, Blood flow and mechanical properties of healing bone, Acta Orthop. Scand. 63 (1992) 487–491. doi:10.3109/17453679209154720.

[12] J. Reeve, M. Arlot, R. Wootton, C. Edouard, M. Tellez, R. Hesp, J.R. Green, P.J. Meunier, Skeletal Blood Flow, Iliac Histomorphometry, and Strontium Kinetics in Osteoporosis: A Relationship Between Blood Flow and Corrected Apposition Rate, J. Clin. Endocrin. & Metab. 66 (1988) 1124–1131. doi:10.1210/jcem-66-6-1124.

[13] G. Kerckhofs, S. Stegen, N. van Gastel, A. Sap, G. Falgayrac, G. Penel, M. Durand, F.P. Luyten, L. Geris, K. Vandamme, T. Parac-Vogt, G. Carmeliet, Simultaneous three-dimensional visualization of mineralized and soft skeletal tissues by a novel microCT contrast agent with polyoxometalate structure, BIomater. 159 (2018) 1–12. doi:10.1016/j.biomaterials.2017.12.016.

[14] O. Barou, S. Mekraldi, L. Vico, G. Boivin, C. Alexandre, M.H. Lafage-Proust, Relationships between trabecular bone remodeling and bone vascularization: a quantitative study, Bone. 30 (2002) 604–612. doi:10.1016/S8756-3282(02)00677-4.

[15] Z. Yao, M.-H. Lafage-Proust, J. Plouët, S. Bloomfield, C. Alexandre, L. Vico, Increase of Both Angiogenesis and Bone Mass in Response to Exercise Depends on VEGF, J Bone Miner Res. 19 (2004) 1471–1480. doi:10.1359/JBMR.040517.

[16] J.D. Boerckel, B.A. Uhrig, N.J. Willett, N. Huebsch, R.E. Guldberg, Mechanical regulation of vascular growth and tissue regeneration in vivo, Proc. Natl. Acad. Sci. USA. 108 (2011) E674–E680. doi:10.1073/pnas.1107019108.

[17] D. Briers, D.D. Duncan, E.R. Hirst, S.J. Kirkpatrick, M. Larsson, W. Steenbergen, T. Stromberg, O.B. Thompson, Laser speckle contrast imaging: theoretical and practical limitations, J. Biomed. Opt. 18 (2013) 066018. doi:10.1117/1.JBO.18.6.066018.

[18] B. Roche, V. David, A. Vanden-Bossche, F. Peyrin, L. Malaval, L. Vico, M.-H. Lafage-Proust, Structure and quantification of microvascularisation within mouse long bones: What and how should we measure?, Bone. 50 (2012) 390–399. doi:10.1016/j.bone.2011.09.051.

[19] J.P. Dyke, R.K. Aaron, Noninvasive methods of measuring bone blood perfusion, Ann. N.Y. Acad. Sci. 1192 (2009) 95–102. doi:10.1111/j.1749-6632.2009.05376.x.

[20] J.T. Au, G. Craig, V. Longo, P. Zanzonico, M. Mason, Y. Fong, P.J. Allen, Gold Nanoparticles Provide Bright Long-Lasting Vascular Contrast for CT Imaging, Am. J. Roentgenol. 200 (2013) 1347–1351. doi:10.2214/AJR.12.8933.

[21] D.P. Clark, K. Ghaghada, E.J. Moding, D.G. Kirsch, C.T. Badea, In vivo characterization of tumor vasculature using iodine and gold nanoparticles and dual energy micro-CT, Phys. Med. Biol. 58 (2013) 1683. doi:10.1088/0031-9155/58/6/1683.

[22] G.E. Nilsson, T. Tenland, P.A. Oberg, Evaluation of a Laser Doppler Flowmeter for Measurement of Tissue Blood Flow, IEEE T. Biomed. Eng. BME-27 (1980) 597–604. doi:10.1109/TBME.1980.326582.

[23] M.F. Swiontkowski, Laser Doppler Flowmetry—Development and Clinical Application, Iowa Orthop. J. 11 (1991) 119–126.

[24] J.F. Griffith, Imaging vasculature of bone, Journal of Orthopaedic Translation. 4 (2014) 192. doi:10.1016/j.jot.2014.07.106.

[25] S. Hellem, L.S. Jacobsson, G.E. Nilsson, D.H. Lewis, Measurement of microvascular blood flow in cancellous bone using laser Doppler flowmetry and 133Xe-clearance, Int. J. Oral Surg. 12 (1983) 165–177. doi:10.1016/S0300-9785(83)80063-5.

[26] P. Okunieff, X. Wang, P. Rubin, J.N. Finkelstein, L.S. Constine, I. Ding, Radiation-induced changes in bone perfusion and angiogenesis, Int. J. Radiat. Oncol. 42 (1998) 885–889. doi:10.1016/S0360-3016(98)00339-3.

[27] M. Melnyk, T. Henke, L. Claes, P. Augat, Revascularisation during fracture healing with soft tissue injury, Arch Orthop Trauma Surg. 128 (2008) 1159–1165. doi:10.1007/s00402-007-0543-0.

[28] B. Roche, A. Vanden-Bossche, M. Normand, L. Malaval, L. Vico, M.-H. Lafage-Proust, Validated Laser Doppler protocol for measurement of mouse bone blood perfusion — Response to age or ovariectomy differs with genetic background, Bone. 55 (2013) 418–426. doi:10.1016/j.bone.2013.03.022.

[29] N.J. Hanne, A.J. Steward, E.D. Easter, S.V. Pinnamaraju, J.H. Jacqueline H, Diet-Induced Obesity Deteriorates Cancellous Bone Structure Despite Increased Blood Perfusion, Orthopaedic Research Society Annual Meeting, San Diego, CA, March 19–22, 2017. Poster #1699.

[30] N.J. Hanne, A.J. Steward, S.V. Pinnamaraju, S.D. Teeter, J.H. Cole, Exercise Therapy Mitigates Reductions in Tibial Blood Flow during Acute Stroke Recovery, Orthopaedic Research Society Annual Meeting, San Diego, CA, March 19–22, 2017. Paper #0337.

[31] M.T. Goova, J. Li, T. Kislinger, W. Qu, et al, Blockade of receptor for advanced blycation end-products restores effective wound healing in diabetic mice, Am. J. Pathol. 159 (2001) 513–25.

[32] H.M. Frost, Bone’s Mechanostat: A 2003 Update, Anat. Rec. 275A (2003) 1081–1101. doi:10.1002/ar.a.10119.

[33] H. Leblond, M. L’Espérance, D. Orsal, S. Rossignol, Treadmill Locomotion in the Intact and Spinal Mouse, J. Neurosci. 23 (2003) 11411–11419.

[34] A.D. Kloos, L.C. Fisher, M.R. Detloff, D.L. Hassenzahl, D.M. Basso, Stepwise motor and all-or-none sensory recovery is associated with nonlinear sparing after incremental spinal cord injury in rats, Experimental Neurology. 191 (2005) 251–265. doi:10.1016/j.expneurol.2004.09.016.

[35] E. Redondo-Castro, A. Torres-Espín, G. García-Alías, X. Navarro, Quantitative assessment of locomotion and interlimb coordination in rats after different spinal cord injuries, J. Neurosci. Methods. 213 (2013) 165–178. doi:10.1016/j.jneumeth.2012.12.024.

[36] S. Amar, Q. Zhou, Y. Shaik-Dasthagirisaheb, S. Leeman, Diet-Induced Obesity in Mice Causes Changes in Immune Responses and Bone Loss Manifested by Bacterial Challenge, Proceedings of the National Academy of Sciences of the United States of America. 104 (2007) 20466–20471.

[37] M. da S. Krause, A. Bittencourt, P.I.H. de Bittencourt, N.H. McClenaghan, P.R. Flatt, C. Murphy, P. Newsholme, Physiological concentrations of interleukin-6 directly promote insulin secretion, signal transduction, nitric oxide release, and redox status in a clonal pancreatic β-cell line and mouse islets, Journal of Endocrinology. 214 (2012) 301–311. doi:10.1530/JOE-12-0223.

[38] K.T. Keylock, V.J. Vieira, M.A. Wallig, L.A. DiPietro, M. Schrementi, J.A. Woods, Exercise accelerates cutaneous wound healing and decreases wound inflammation in aged mice, A. J. Physiol.-Reg. I. 294 (2008) R179–R184. doi:10.1152/ajpregu.00177.2007.

[39] V.L. Ferguson, R.A. Ayers, T.A. Bateman, S.J. Simske, Bone development and age-related bone loss in male C57BL/6J mice, Bone. 33 (2003) 387–398. doi:10.1016/S8756-3282(03)00199-6.

[40] J.M. Somerville, R.M. Aspden, K.E. Armour, K.J. Armour, D.M. Reid, Growth of C57BL/6 mice and the material and mechanical properties of cortical bone from the tibia, Calcif. Tissue Int. 74 (2004) 469–475. doi:10.1007/s00223-003-0101-x.

[41] S.C. Marks, P.R. Odgren, Chapter 1 - Structure and Development of the Skeleton, in: J.P. Bilezikian, L.G. Raisz, G.A. Rodan (Eds.), Principles of Bone Biology (Second Edition), Academic Press, San Diego, 2002: pp. 3–15. doi:10.1016/B978-012098652-1.50103-7.

[42] H.-P. Gerber, T.H. Vu, A.M. Ryan, J. Kowalski, Z. Werb, N. Ferrara, VEGF couples hypertrophic cartilage remodeling, ossification and angiogenesis during endochondral bone formation, Nat Med. 5 (1999) 623–628. doi:10.1038/9467.

[43] J.M. Wallace, R.M. Rajachar, M.R. Allen, S.A. Bloomfield, P.G. Robey, M.F. Young, D.H. Kohn, Exercise-induced changes in the cortical bone of growing mice are bone- and gender-specific, Bone. 40 (2007) 1120–1127. doi:10.1016/j.bone.2006.12.002.

[44] J. Prasad, B.P. Wiater, S.E. Nork, S.D. Bain, T.S. Gross, Characterizing gait induced normal strains in a murine tibia cortical bone defect model, J. Biomech. 43 (2010) 2765–2770. doi:10.1016/j.jbiomech.2010.06.030.

[45] A.G. Berman, M.J. Hinton, J.M. Wallace, Treadmill running and targeted tibial loading differentially improve bone mass in mice, Bone Rep. 10 (2019). doi:10.1016/j.bonr.2019.100195.

[46] R. Ellman, J. Spatz, A. Cloutier, R. Palme, B.A. Christiansen, M.L. Bouxsein, Partial reductions in mechanical loading yield proportional changes in bone density, bone architecture, and muscle mass, J. Bone. Miner. Res. 28 (2013) 875–885. doi:10.1002/jbmr.1814.

[47] A.J. Steward, J.H. Cole, F.S. Ligler, E. Loboa, Mechanical and Vascular Cues Synergistically Enhance Osteogenesis in Human Mesenchymal Stem Cells, Tissue Eng. Part A. (2016). doi:10.1089/ten.TEA.2015.0533.

[48] D.E. Hurwitz, D.R. Sumner, T.P. Andriacchi, D.A. Sugar, Dynamic knee loads during gait predict proximal tibial bone distribution, J. Biomech. 31 (1998) 423–430. doi:10.1016/S0021-9290(98)00028-1.

[49] K. Gertz, J. Priller, G. Kronenberg, K.B. Fink, B. Winter, H. Schröck, S. Ji, M. Milosevic, C. Harms, M. Böhm, U. Dirnagl, U. Laufs, M. Endres, Physical Activity Improves Long-Term Stroke Outcome via Endothelial Nitric Oxide Synthase—Dependent Augmentation of Neovascularization and Cerebral Blood Flow, Circ. Res. 99 (2006) 1132–1140. doi:10.1161/01.RES.0000250175.14861.77.

[50] M. Styner, G.M. Pagnotti, C. McGrath, X. Wu, B. Sen, G. Uzer, Z. Xie, X. Zong, M.A. Styner, C.T. Rubin, J. Rubin, Exercise Decreases Marrow Adipose Tissue Through β-Oxidation in Obese Running Mice, J Bone Miner Res. 32 (2017) 1692–1702. doi:10.1002/jbmr.3159.

[51] L.E. Kuo, Z. Zukowska, Stress, NPY and vascular remodeling: Implications for stress-related diseases, Peptides. 28 (2007) 435–440. doi:10.1016/j.peptides.2006.08.035.

[52] C. Colnot, X. Zhang, M.L.K. Tate, Current insights on the regenerative potential of the periosteum: Molecular, cellular, and endogenous engineering approaches, J. Orthop. Res. 30 (2012) 1869–1878. doi:10.1002/jor.22181.

[53] M.J. Kowalski, E.H. Schemitsch, P.J. Kregor, D. Senft, M.F. Swiontkowski, Effect of periosteal stripping on cortical bone perfusion: A laser doppler study in sheep, Calcif. Tissue Int. 59 (1996) 24–26. doi:10.1007/s002239900080.

[54] R.E. Tomlinson, M.J. Silva, K.I. Shoghi, Quantification of Skeletal Blood Flow and Fluoride Metabolism in Rats using PET in a Pre-Clinical Stress Fracture Model, Mol. Imaging Biol. 14 (2012) 348–354. doi:10.1007/s11307-011-0505-3.

